# Microbial Ecosystems Reveal a Universal Signature of Ecological Assembly

**DOI:** 10.64898/2026.07.10.737833

**Authors:** James Holehouse, Christopher P. Kempes, Geoffrey B. West, Anshuman Swain

## Abstract

Microbial communities obey universal macroecological scaling laws, but which sub-processes generate them remains debated across competing theoretical frameworks. Here we calibrate a modified Yule-Simon model to metagenomic data from 11 distinct microbial environments, revealing that microbial ecosystems occupy a qualitatively distinct region of a two-parameter mechanistic space, characterized by near-neutral recruitment (i.e., linear preferential attachment) and broad diversification strategies, unlike any previously studied complex system, including prokaryotic proteomes and urban economies. This distinctive position, confirmed analytically and validated against sparse-data robustness tests, provides both a mechanistic explanation for observed self-similarities in microbial rank-frequency distributions and a quantitative signature of ecological assembly.

---

Microbial communities exhibit robust macroecological scaling laws across environments spanning orders of magnitude in size: the number of distinct species scales sublinearly with community size following Heaps’ law [1, 2], species rank-frequency distributions approach near-universal functional forms [1], and abundance fluctuations exhibit consistent statistical structure [1]. Existing theoretical frameworks each describe part of this picture, but most characterize the converged distribution rather than decomposing the sub-processes that generate it. Hubbell’s Unified Neutral Theory of Biodiversity (UNTB) [3] reproduces log-series abundance distributions but rests on the empirically contested assumption of ecological equivalence [4]. The Maximum Entropy Theory of Ecology [5] predicts macroecological patterns from macroscopic state variables at equilibrium; its dynamic extension, DynaMETE [6], incorporates disturbance but does not decompose origination from recruitment. Niche- and competition-based theories [7] instead emphasize deterministic species sorting over stochastic assembly. We therefore lack a parsimonious, mechanistically interpretable model that simultaneously explains diversity scaling and rank-frequency structure across multiple environments. Understanding what generates these universal patterns has become a central challenge in (microbial) ecology, with direct implications for predicting community responses to environmental perturbation, interpreting the rapidly expanding archive of metagenomic surveys, and designing microbiome-based interventions [2, 8].

We address this gap by applying a modified Yule-Simon generative model [9, 10] to metagenomic data from 11 distinct microbial environments drawn from the comprehensive dataset of Grilli [1, 11], originally obtained from EBI Metagenomics [12]. The model describes community assembly as a stochastic process in which each new individual either establishes a new species (with probability *p*) or joins an existing one (with probability 1 − *p*). It is governed by two ecologically interpretable parameters: a diversification feedback exponent *θ*, which determines how the per-individual probability of new-species establishment varies with community composition (*θ <* 0 corresponds to facilitation; *θ >* 0 to competitive suppression), and a preferential attachment exponent *γ*, which governs abundance-dependent recruitment (*γ* = 1 recovers neutral proportional recruitment; *γ >* 1 amplifies dominance). Specifically, this parametrization is given by

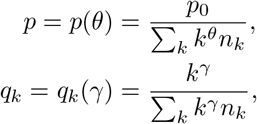

wherein *p*_0_ is a multiplicative constant, *q*_*k*_ is the probability of expanding a function of size *k*, and *n*_*k*_ is the number of species with abundance *k*. For more details refer to the *Online Methods*. This parameterization generalizes the classical Yule-Simon process [13, 14] and was recently shown to quantitatively explain functional diversity scaling in prokaryotic proteomes, U.S. federal agencies, and urban economies [9, 10]. Our goal is to locate microbial ecosystems within this shared mechanistic space.

Across complex adaptive systems, an important empirical law is Heaps’ law, which states that the number of unique system functions—the diversity, denoted *D*(*N*)—scales as a power law in the system size, i.e., *D*(*N*) ∼ *N* ^*β*^ [9, 10, 15]. For the microbial ecosystems studied herein, Heaps’ law is confirmed across all 11 environments (Table S1, Fig. S1), with scaling exponents *β* ranging from 0.24 (oral cavity) to 0.66 (aquatic marine), all significantly different from zero. The consistent sublinear scaling (*β <* 1) reflects a universal tendency for new-species discovery to decelerate as communities grow, while the substantial variation in *β* motivates a mechanistic rather than purely phenomenological characterization. Fitting the model to the 10 most species-rich communities per environment via likelihood-free inference, we recover excellent fits to empirical rank-frequency distributions across all environments and three orders of magnitude in ecosystem size (Fig. 2, insets; Figs. S3 and S4).

**FIG. 1:**
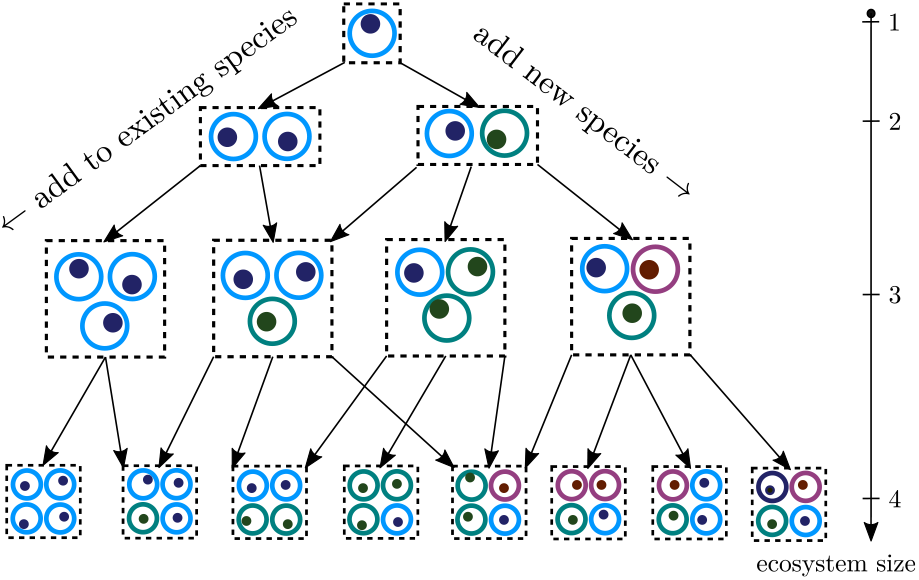
Illustration of the mechanistic model of ecological assembly, adapted from [9]. There are two key mechanistic processes: (i) adding a new species, parameterized by *θ*; and (ii) expanding existing species with a generalized preferential attachment mechanism, parameterized by *γ*. For more details, please refer to the *Model* Section in the *Online Methods*.

**FIG. 2:**
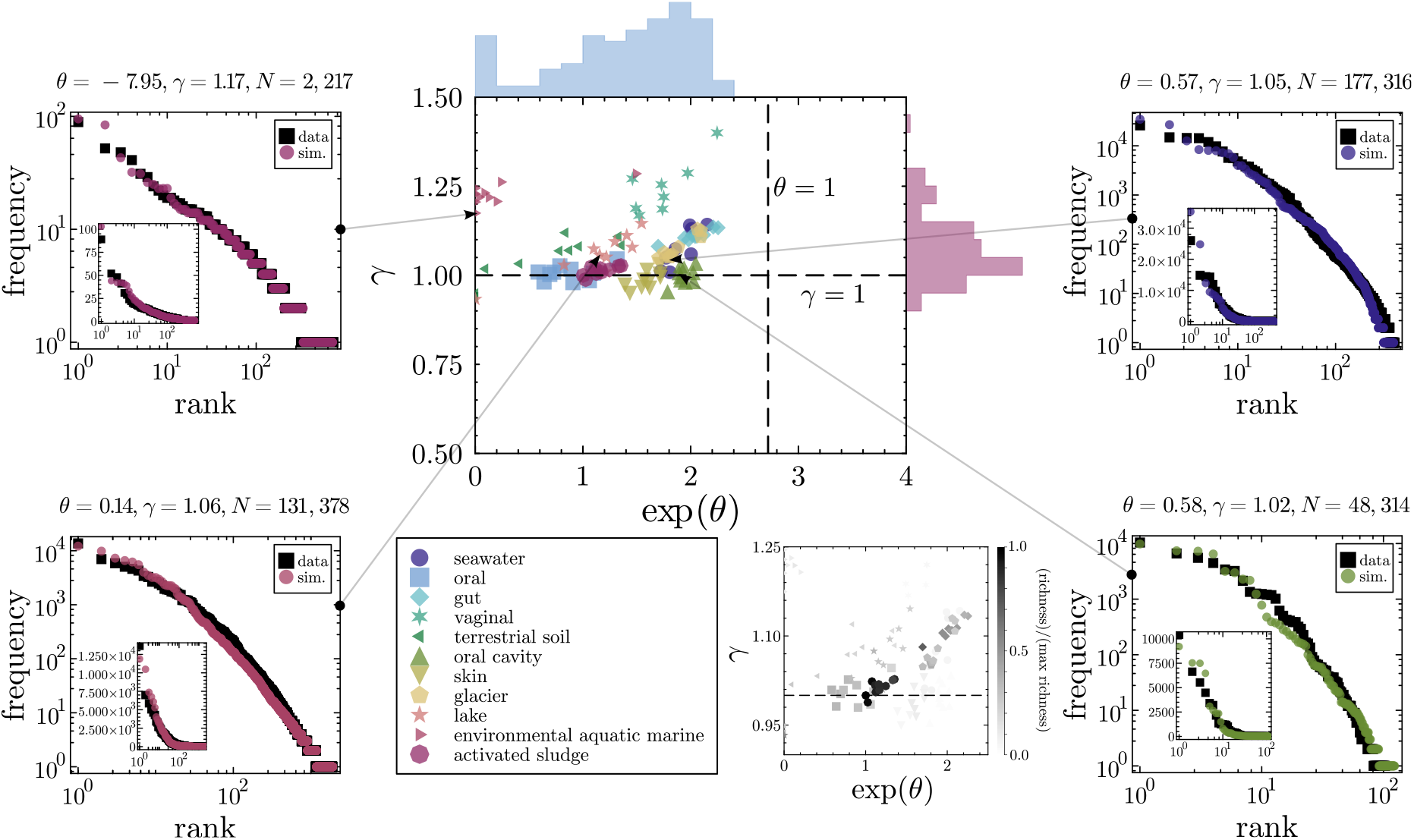
Calibration of the generative model to microbial metagenomic data across 11 environments. The central phase diagram on the top row shows the positions of microbial ecosystems from 11 distinct environments in the space of phenomenological parameters *θ* and *γ*. For each environment, the top 10 ecosystems as measured by number of reads were fit to the model; details of the fitting procedure are provided in the *Online Methods*. The *θ*-axis is exponentiated due to the prevalence of negative values in the *environmental aquatic marine* environment. Histograms on the axes show the marginal spread of *γ* and *θ* across all ecosystems; mean values are *γ*_mean_ = 1.07 (median 1.06) and *θ*_mean_ = −0.37 (median 0.42). While the modal value of *γ* indicates that many ecosystems exhibit linear preferential attachment at *γ* ≈ 1, the spread in the histograms shows that ecosystems range from strongly negative *θ* (facilitation) to *θ* approaching 1 (competitive suppression), reflecting a wide diversity of ecological assembly mechanisms. Strong super-linear preferential attachment (*γ >* 1.2) is observed only in smaller ecosystems (*environmental aquatic marine* and *vaginal*, with typical sizes of order *O*(10^4^) and *O*(10^3^) reads respectively), suggesting that such dynamics are not viable in larger communities. Four example model fits are shown with insets confirming agreement in linear-log rank-frequency space. The bottom row re-plots the phase diagram coloured by species richness (the richest ecosystem has 8,653 species, in *activated sludge*), emphasizing that the most species-rich ecosystems cluster near *γ* ≈ 1.

The resulting phase diagram (Fig. 2) reveals a coherent structure with two striking features. First, the preferential attachment exponent clusters tightly near the neutral boundary, with *γ*_mean_ = 1.07 (median 1.06), indicating that recruitment is, to first approximation, proportional to species abundance across all 11 environments. Second, while *γ* is nearly universal, the diversification feedback spans a much wider range—from strongly negative values (aquatic marine, indicating rapid facilitated diversification) to values approaching 1 (gut, skin), with mean *θ*_mean_ = ™0.37 (median 0.42). This asymmetry reveals that the mechanism of recruitment is broadly shared among microbial communities, while the mechanism governing diversification rate varies substantially across ecosystem types. Strong superlinear preferential attachment (*γ >* 1.2) is confined to the smallest ecosystems (vaginal, *N* ∼ 10^3^; aquatic marine, *N* ∼ 10^4^). Two mechanisms likely contribute: community size may be insufficient for the condensate formation predicted analytically for *γ >* 1 to manifest at finite *N* [16], and downsampling systematically inflates inferred *γ* in the *γ >* 1 regime (see *Online Methods*).

This position in parameter space is qualitatively distinct from all previously studied complex systems within this framework. Prokaryotic proteomes, federal agencies, and cities all exhibit sublinear preferential attachment (*γ <* 1); cities in particular sit near *θ* ≈ 1, reflecting strong competitive suppression of new specializations [10]. Microbial ecosystems occupy the orthogonal corner: *γ* ≈ 1, *θ* ∈ [−1, 1]—a combination not observed in any previously studied system. This finding also challenges the longstanding sociological analogy between cities and ecological communities [17]: in this parameter space, cities sit near organisms rather than near ecosystems, consistent with the view that urban functional diversity grows through suppression-dominated optimization-like processes rather than facilitated ecological assembly [18]. The self-similarities in rank-frequency distributions observed across cities and ecosystems have qualitatively distinct origins: in cities, self-similarity is diversity-driven, requiring both *γ <* 1 and logarithmic diversity scaling [9]; in microbial ecosystems, it is preferential-attachment-driven—any system with *γ* = 1 exhibits self-similar rank-frequency distributions independently of diversity scaling, as predicted analytically. This shows that identical rank-frequency curves can arise from distinct generative processes—near-neutral, MaxEnt-derived, or niche-based—so discriminating between them requires fitting the full (*θ, γ*) pair rather than comparing distributional fit alone. A practical corollary is that raw rank-frequency curves directly report *γ*: collapse across differently sized communities—as seen in oral, oral cavity, and activated sludge (Fig. S2)—is direct, model-free evidence for *γ* ≈ 1.

The prevalence of *γ* ≈ 1 has a natural ecological interpretation: linear preferential attachment is formally equivalent to the zero-sum sampling of Hubbell’s UNTB [3], but our framework infers this effective neutrality from data rather than assuming it a priori. This matters mechanistically: whereas the UNTB posits equivalence as a starting assumption, our model treats *γ* as a free parameter and recovers near-neutrality as an emergent property of the most species-rich environments. The broad distribution of *θ* represents a range of diversification strategies across distinct environments. The most novel regime, *θ <* 0, represents facilitated diversification—likely reflecting the tendency of diverse microbial communities to generate metabolic byproducts, establish syntrophic and cross-feeding networks, and condition local environments in ways that lower colonization barriers for additional species, creating a positive feedback between existing diversity and future diversifiability. This contrasts fundamentally with other ecosystems, but notably all nonecological complex systems studied in this framework, where *θ >* 0 indicates that existing functions always suppress new ones. This suggests that facilitation can be a structural property of mature microbial communities rather than just a transient feature of early assembly. This is notable in light of diversity-dependence theory in macroevolution, where high existing species richness is thought to dampen speciation rates; in microbial ecosystems, high diversity appears instead to accelerate further diversification, inverting the macroevolutionary expectation.

An important practical consequence of the *γ* ≈ 1 regime concerns data sparsity. Metagenomic reads are a partial, downsampled observation of the underlying community [8, 19]. Robustness analysis (*Online Methods*, Fig. S6) demonstrates that in the *γ* ≈ 1 regime, downsampled communities are nearly statistically indistinguishable from their fully realized counterparts—a direct consequence of the analytical self-similarity of the *γ* = 1 solution. Parameter inference is therefore most reliable precisely where the majority of microbial ecosystems are located, validating the use of metagenomic data for mechanistic inference. This contrasts with the *γ <* 1 regime, characteristic of proteins and federal agencies, where downsampling underestimates both *γ* and *θ*, making inference unreliable without correction. The concentration of microbial ecosystems near *γ* ≈ 1 therefore implies not only that they are amenable to mechanistic inference from sparse data, but that the near-neutral attractor may itself confer compositional stability, since self-similar rank-frequency structure is inherently more reproducible and predictable. This connects to the emerging concept of emergent predictability [20]: as richness grows, coarse compositional descriptions become more predictive of community function, consistent with communities converging on the analytically tractable *γ* = 1 attractor.

In summary, calibrating a two-parameter generative model to metagenomic data from 11 microbial environments reveals that ecological community assembly occupies a precise, quantitative address in the space of complex-systems mechanisms: a range of diversity generating strategies—from facilitated diversification (*θ <* 0) to competitive suppression (*θ >* 0)—acting in concert with near-neutral recruitment (*γ* ≈ 1). This combination is not observed in any previously characterized biological or socioeconomic system, suggesting that facilitation and near-neutrality are not incidental properties of microbiomes but the mechanistic signature of ecological assembly itself. The framework simultaneously accounts for Heaps’ law exponents, rank-frequency self-similarity, and the robustness of metagenomic inference within a single two-parameter model, and provides a quantitative basis for cross-system comparison. Applying the same calibration to macro-organismal communities—animal gut systems, soil metacommunities, plant–pollinator networks—will directly test whether the ecological assembly signature identified here is universal to life or specific to the microbial scale.

## ACKNOWLEDGMENTS

JH and AS started this project based on discussion at the Santa Fe Institute’s 2025 *Postdocs in Complexity* Conference. JH would like to thank Paul Krapivsky and Harrison Hartle for useful discussions. We would like to thank the UNM Center for Advanced Research Computing, supported in part by the National Science Foundation, for providing the research computing resources used in this work.

## AUTHOR CONTRIBUTIONS

JH and AS conceived the manuscript, with input from CK and GBW. JH and AS wrote the first draft of the paper. All authors revised and edited the paper. CK and AS sourced the data used in the study. JH performed the analytic and computational analyses. JH made the figures.

## AI USAGE STATEMENT

All first drafts of writing were performed by human researchers. AI was used to find grammatical and typographical mistakes at the end of the writing process. Generative AI (ChatGPT) was used to aid the derivation of analytics in the regime of *γ >* 1.

## DATA AND CODE AVAILABILITY

All data and code used can be found at: https://github.com/jamesholehouse/Ecological-Preferential-Attachment.

## ONLINE METHODS

### Model

We introduce a model of organization growth that was previously used to describe the evolution of diversity of protein abundances in prokaryotes and occupations within cities and federal agencies (see refs. [9, 10]). The model is a modified form of Yule-Simon process [13, 14] in which as an organization grows, new elements are either added to an *existing function*, or they create their own *new function*. In an ecological context, an existing *function* is a *species* that is already present in the ecosystem, while a *new function* represents the introduction of a *new species*—either by speciation or migration. The core mathematical object is the number of functions of size *k* for a given ecosystem size *N* , which we denote as *n*_*k*_ = *n*_*k*_(*N*), and it satisfies the dynamical equation

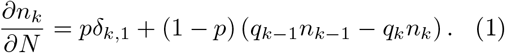

where ∂*n*_*k*_*/*∂*N ≈ n*_*k*_(*N* + 1) − *n*_*k*_(*N*), *p* is the probability of adding a new species at size *N* , and the *q*_*k*_ encode the probabilities of attaching to existing functions of size *k*. From *n*_*k*_, the rank-frequency and normalized rank-frequency distributions can be easily derived [9, 21]. Normalized by *N* , *n*_*k*_ becomes the abundance distribution *ρ*_*k*_ = *n*_*k*_*/N*.

As the ecosystem grows to its required size, the number of unique species (also known as the diversity or richness) will increase as new species are added. Mathematically, the diversity, denoted *D*(*N*), can be represented by either

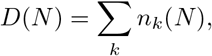

or

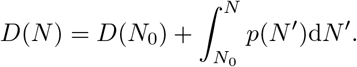

The first of these two definitions states that the number of unique species is the sum over the number of species of each given size *k*. The second definition states that the number of unique species is the integral over the probability of adding a new species at each step, plus the original number *D*(*N*_0_). Additionally, existing species will grow in size based on a preferential attachment mechanism, specified by *q*_*k*_. Note that the total number of organisms in the system is self-consistently given by *N* = ∑ _*k*_ *kn*_*k*_.

Because the data provides the diversity of species as the number of unique metagenomic reads, given a specific number of *reads* (and not organisms), one can think of each time a species is added to the ecosystem as corresponding to a read being measured in the metagenomic sequencing. We return to this point in the Sections *Data* and *Robustness check*.

We parameterize *p* and *q*_*k*_ as follows:

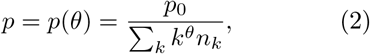

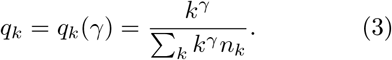

The parameter *θ* determines the probability *p*(*θ*) = *p*(*θ, N*) of adding new species (a decreasing function of *N*), *p*_0_ is a scale factor of *p* which does not play a phenomenological role, and *γ* is the strength of preferential attachment to add members to existing species. While *q*_*k*_ is the standard extension of the linear preferential attachment rule (*γ* = 1) to generalized preferential attachment rule [22], the form of *p* is chosen because: (i) this parameterization can capture a substantial range of diversity growth curves across many complex systems [10]; (ii) from single parameter (*θ*) we get a highly flexible functional form of *p*; and (iii) its similar functional form to the preferential attachment rule allows for a unified mathematical treatment, and the prediction of empirically observed self-similarities in data [9]. Later, we show how to solve the model in ecologically realistic regimes wherein *γ* ≥ 1.

While this introduction to the model takes a largely mathematical approach, the easiest way to use the model is to simulate it. Analytic difficulties that arise due to the model’s considerable flexibility—i.e., the free parameters *θ* and *γ*—mean that generally only asymptotic results are possible (in the regime of large *N* , see *Analytics of the model*). To calibrate the data to the model we simulate the model and infer the appropriate parameters by minimizing a distance function between the data and the model’s prediction. This simulation algorithm follows the standard stochastic simulation algorithm (SSA) approach common in models of chemical kinetics [23].

### Data

We utilize data popularized in a seminal article of Grilli [1, 11], who obtained the data from EBI Metagenomics [12]. The data consists of 14 different microbial environments (e.g., *sea-water, oral*, etc.), and within each environment there are many samples of different microbial ecosystems. In our analyses, three of the environments had to be discarded: (i) the *feces* environment consisted of only three samples, which is too few samples to to reliably fit a diversity scaling curve (as seen in Fig. S1); (ii) both the *marine hydrothermal vents* and *river* environments had diversity scaling exponents *β* that were not significantly different than zero (see Table I in the SI), due to data sparsity and limited ecosystem size ranges respectively. Since our model predicts that microbial diversity should increase with ecosystem size/number of metagenomic reads, these two environments were also discarded from our analyses.

This left us with 11 environments to calibrate our model against, whose environment labels are shown in Fig. 2. These are wide ranging, from human microbiomes to glaciers to activated sludge, and individual ecosystems utilized in our study span four orders of magnitude, from *N* ∼ *O*(10^3^) to *N* ∼ *O*(10^6^). Note that from this data, we assumed that the total number of reads is reflective of the size of a community for a given diversity of species, even though this observation is partial. In Section *Robustness check*, we assess the robustness of this assumption by inferring the parameters of down-sampled simulated data for which *θ* and *γ* are known parameters. This check allows us to understand the implications of parameter inference on data that is sparser than the real ecosystems that data describes, and to observe systematic parameter drift in different regions of this space.

### Calibration procedure

We use a likelihood-free inference algorithm [10] to fit the parameters *θ* and *γ* to the 10 most populous ecosystems in each environment (also see ref. [10, SI Sec. 4B]). In each case, *p*_0_ was predicted from the scaling of the diversity curves using a method developed in ref. [10, SI Sec. 4D]. This meant that inference of only the most important phenomenological parameters *θ, γ* was necessary. In what follows, we summarize the steps of the calibration procedure.

First, for each ecosystem we defined the boundaries of the parameter space, which were chosen as *θ* ∈ [−10, 1.5] and *γ* ∈ [0.2, 1.5]. These values were selected such that calibrated parameter values were not found at the boundaries of the parameter space. Note that in some cases the lower bound of the *θ* axis of the parameter space was reached (mainly in the *environmental aquatic marine* environment)—indicating that for the environments with the steepest Heaps’ law exponents that the model’s abilities to capture the data were limited in some cases. Additionally, we specified an initial condition for each ecosystem that consisted of the (second) smallest ecosystem found in that particular environment. These initial conditions are shown in the first column of Figs. S3 and S4.

Second, we used an adaptive differential evolution algorithm [24] (implemented via the Julia package `BlackBoxOptim` [25]) to sample the parameter space and minimize the distance between the rank-frequency distributions from the data and those predicted from the model. The distance metric used was the sum of the logarithm of the difference between species of the same rank in the simulated versus empirical data. This cost function is sensitive enough that the resultant rank-frequency distributions are well fit even in linear space (see insets of calibration results in Fig. 2).

Third, optimization was continued until a specified time limit of 44 hours was reached. This allowed even the largest of the ecosystems enough time to converge on an optimum. Calibrations were performed on the University of New Mexico’s CARC supercomputer, with each system run on a CPU equipped with 5 GB of RAM. The results of all the calibrations are shown in Fig. 2, and the rank-frequency distributions of the three largest ecosystems in each environment are shown in the final three columns of Figs. S3 and S4.

### Analytics of the model

*The γ* = 1 *regime*. This regime of the model was asymptotically solved in ref. [9], although the systems studied therein did not admit linear preferential attachment. For microbial ecologies, linear preferential attachment is a common trait—as seen from our calibrations in Fig. 2—and therefore we re-derive the solution here for comparison to the data.

The first step of the solution involves solving Eq. (1) in the regime of *N* ≫ 1 and where 1 − *p* ≈ 1. Noting that ∑_*k*_ *kn*_*k*_ = *N* and solving for *n*_1_ reveals ∂*n*_1_(*N*) ∼ *p*. One can show that in the regime of *γ* ≤ 1 that [9, Sec. S2]

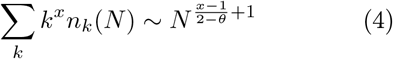

meaning that *p ∼ N* ^−*ν*^ with *ν* = 1*/*(2 − *θ*). The sanity check ∑_*k*_ *kn*_*k*_ = *N* validates the scaling of the sum. Using Eq. (4) results in *n*_1_(*N*) ∼ *N* ^1−*ν*^. As an ansatz, let *n*_*k*_ ∼ *a*_*k*_*N* ^1−*ν*^. To find the *a*_*k*_, we substitute the ansatz for *n*_*k*_ into Eq. (1). This results in

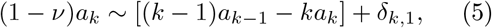

where conveniently the *N* dependence on each side of the equation cancels out. Solving this two-term recurrence relation gives the asymptotic solution

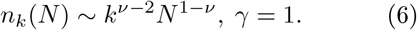

To find the rank *r* of a species with abundance *k*, consider the definition

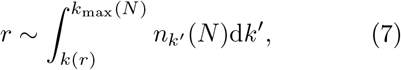

where in *k*_max_(*N*) is the size of the maximum function, which in the regime of linear preferential attachment scales proportionately with *N* . Evaluating the integral gives the following scaling of the rank-frequency distribution,

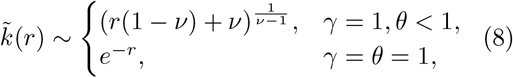

which notably is independent of *N* , indicating the self-similarity (i.e., size independence) of the rank-frequency curves when *γ* = 1.

Another useful representation of an organization is the normalized rank-frequency distribution, wherein the frequency is a function of the normalized rank 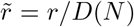. The utility of the normalized rank-frequency distribution is that it allows one to see how the abundance is spread across functions in a way that is independent of the diversity. Substituting 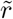 into Eq. (8) and utilizing the Jacobian 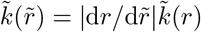 gives

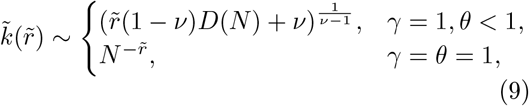

where we see that the self-similarity observed for the rank-frequency distribution vanishes for the normalized rank-frequency distribution.

In summary, in the regime of *γ* = 1:

1. Rank-frequency distributions show self-similar behaviors that are independent of *N*.
2. Rank-frequency distributions are Zipfian for *θ <* 1.
3. Normalized rank-frequency distributions do not show self-similar behaviors.

*The γ >* 1 *regime*. In this regime the scaling of the sum in Eq. (4) breaks down. One can show this as the probability of joining a function of size *k, q*_*k*_ ∼ *k*^*γ*^*/*∑_*k*_ *k*^*γ*^*n*_*k*_, would predict the following under the scaling predicted by Eq. (4)

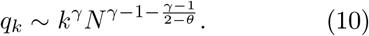

If *θ <* 1 and there exists a species with size scaling *k* = *K*(*N*) ∼ *cN* —which is generally true for super-linear mechanisms of preferential attachment [26]—then this predicts that the probability *q*_*k*_ → ∞ in the asymptotic limit of *N* , which is, of course, impossible.

To make progress, following the literature on preferential attachment in growing random networks [16, 22, 26], we posit that when *γ >* 1 that the largest species grows in proportion with the size of the system, i.e., *k*_max_ = *K*(*N*) ∼ *N* − *O*(*D*(*N*)). This is readily verified from simulations (see Figs. S5(a) and (e)). In the sum scaling this leads to

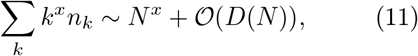

where notably the largest species has *n*_*K*_ = 1. Ecologically speaking, the formation of a *condensate* means that in the regime of super-linear preferential attachment there will be a single species that dominates the ecosystem for *N* ≫ 1.

As in the *γ* = 1 case, we want to solve Eq. (1) in the regime of *N* ≫ 1 and where 1 − *p* ≈ 1. To begin we start by attempting a solution in the *k* ≥ 2 regime, in which *p* does not play a role. Unlike the *γ* = 1 case, an ansatz of the form *n*_*k*_ = *c*_*k*_*N* ^*α*^ does not lead to a balanced asymptotic equation as observed in Eq. (5). Following the literature on growing random networks (e.g., [26, Eq. (12)]), we consider that in the superlinear regime that each *n*_*k*_ has its own asymptotic scaling exponent with *N* , i.e., 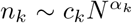. Substituting this into Eq. (1) and using ∑_*k*_ *k*^*γ*^*n*_*k*_ gives

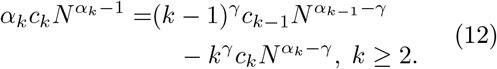

Employing dominant balance [16, 27], for *N* ≫ 1 we see that the second term on the right-hand side of the equation will vanish because *α*_*k*_ − 1 *> α* − *γ*. The resulting equation leads to two separate recurrence relations in *α*_*k*_ and *c*_*k*_. First,

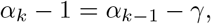

whose solution is *α*_*k*_ = *α*_1_ − (*k* − 1)(*γ* − 1) in which *α*_1_ is to be determined by the boundary condition at *k* = 1. Second, for the *c*_*k*_ we find

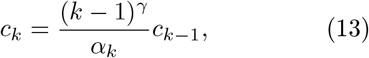

whose solution

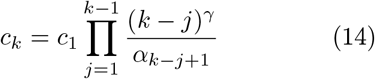

depends on knowledge of both *c*_1_ and *α*_1_.

A few comments are in order. First, the scaling 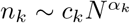 does not include the condensate, it only includes the species of small sizes. We will refer to these species as existing in a *low* -*k cloud*. Second, in this low-*k* cloud, we find that each *n*_*k*_ decays faster with increasing *k*. This indicates that the majority of small species are of size 1 in the limit *N* ≫ 1 (as verified via simulations in Fig. S5(a) and (e)). Finally, Eq. (13) predicts that there is a finite *k* = *k*^⋆^ cutoff, where the *c*_*k*_ would otherwise become negative. The analytic prediction is then that *n*_*k*_ = 0 for *k*^⋆^ *< k < K*. The cutoff *k*^⋆^ is given by,

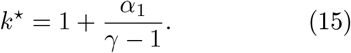

All together, this gives

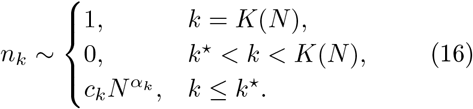

To find *c*_1_ and *α*_1_ we need to find the scaling of *n*_1_. For *k* = 1, we have 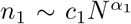, and we find the dominant balance equation

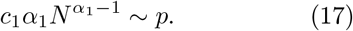

Therefore, understanding the asymptotic scaling of *p* will allow us to find both *α*_1_ and *c*_1_. The asymptotic scaling of *p* relies on the asymptotic scaling of ∑_*k*_ *k*^*θ*^*n*_*k*_. The scaling of this sum can be broken up into two parts (as in Eq. 11). First, there is the contribution coming from the condensate, given by *N* ^*θ*^. Second, there is the contribution coming from the small values of *k*, which is dominated by the number of species of size 1. Together, this gives the scaling

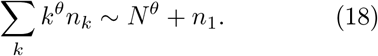

Therefore, following Eq. (17) we find

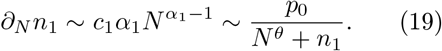

The values of *c*_1_ and *α*_1_ therefore depend on which of *N* ^*θ*^ or *n*_1_ dominate in the denominator of the right-hand side of the equation. First consider that *n*_1_ dominates, i.e., *n*_1_ ≫ *N* ^*θ*^. This results in

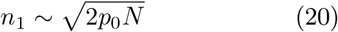

and therefore 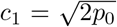 and *α*_1_ = 1*/*2. For the case in which *N* ^*θ*^ ≫ *n*_1_, occurring for *θ >* 1*/*2, we instead find that

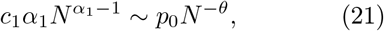

where now one can identify *α*_1_ = 1 − *θ* and *c*_1_ = *p*_0_*/*(1 − *θ*). In sum, this gives

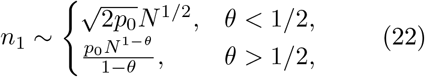

from which *c*_1_ and *α*_1_ are readily identified.

These results also lead to the following diversity scaling which exhibits a second-order phase transition in *θ*

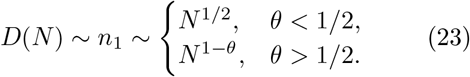

A similar phase transition is not observed for *γ* ≤ 1.

For the rank-frequency distribution, following Eq. (7), we find that the rank, *r*, of a species with *k* members is given by

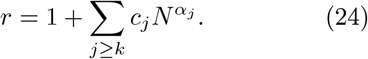

Clearly the condensate retains rank 1, with the species in the small-*k* cloud described by the sum on the right-hand side. Because *α*_*k*_ decreases with respect to *k*, asymptotically one can state that

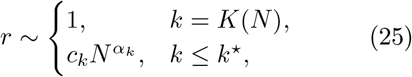

i.e., for *k* ≤ *k*^⋆^ only the largest term in the sum from Eq. (24) is retained. For *γ >* 1 this describes a rank-frequency distribution dominated by a single rank-1 function, with a stepped low abundance curve dominated by functions of size 1. In Fig. S5 we confirm the analytic results predicted from this section for the super-linear regime.

In summary, in the regime of *γ >* 1:

1. A condensate asymptotically forms, indicating that a single species dominates the system’s abundance.
2. Each individual *n*_*k*_ now has its own scaling with system size, in a departure from the consistent scaling observed in the linear and sub-linear preferential attachment cases [9].
3. The scaling of the diversity predicts a discontinuity at *θ* = 1*/*2.
4. The rank-frequency distribution is asymptotically dominated by a single rank-1 species, with a step-like low abundance curve exhibiting a finite cutoff at *k* = *k*^⋆^.

### Robustness check: Model calibration with sparse ecosystems

An important aspect of metagenomic data is that ecosystem diversity is approximated by unique metagenomic reads and ecosystem size is approximated by the number of reads [8, 19]. This type of data is good for assessing relative abundances, but is considered relatively poor in assessing the importance of ecosystem size in the generation of diversity. In reality, if the data show that 10 reads corresponded to a specific species, and there were 1000 unique species found, that could represent a *down-sampling* of the real ecosystem with hundreds of organisms of that species and tens of thousands of unique species. Sampling issues exist in ecology at all scales, not only in metagenomic data, and even for macroscopic organisms such as fish or birds, observed distributions of species are only down-samplings of an ecosystem’s real abundance distributions [28]. This means that we should exercise caution in the mechanistic inferences provided in the main text.

Our modeling framework provides a perfect training ground to test the robustness of the calibrations of *θ* and *γ* on partial, incomplete, or sparse data, and to observe parameter drift between assigned parameter and those inferred from the down-sampled data (via the calibration procedure specified above). Additionally, this allows us to determine the robustness of our calibrations in the main text (see Fig. 2) and to see which areas of the parameter space lead to robust inference—even on necessarily sparse metagenomic data. The fundamental reason for *parameter drift* is that a *simulated ecosystem’s* diversity and organization with assigned parameters *θ*^⋆^ and *γ*^⋆^, say of size *N* = 10^6^, when *down-sampled* to *N* = 10^5^, in general, will not admit the same statistics as an ecosystem *back-tracked* to size *N* = 10^5^ with the same parameters *θ*^⋆^ and *γ*^⋆^ (see Fig. S6(a)–(c)). In the box below, we define key terms and variables used in this section.

In our robustness analysis we take 20 randomly generated parameter sets from the space *γ* ∈ [0.95, 1.2] and *θ* ∈ [− 1, 1], which represents the empirically observed space of parameters from our calibrations on the empirical data (see Fig. 2, histograms on the axes of the phase diagram). Arbitrarily, we set *p*_0_ = 1. For each parameter set, denote *θ*^⋆^ and *γ*^⋆^ as the true parameters. For each parameter set, we simulate an ecosystem of size *N* = 10^6^, which we then systematically down-sample. In the down-sampling procedure, we assign each species a probability equal to its relative abundance in the simulated ecosystem. Then, with those probabilities, we sample *N*_red_ = 10^5^ times, generating a down-sampled ecosystem. Finally, we use the calibration algorithm to infer the parameters *θ*_red_ and *γ*_red_ that best fit the down-sampled data. The values of

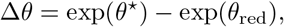

and

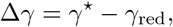

then tell us how parameters drift due to down-sampling.

For these calibrations we used a time limit of 1 day and used University of New Mexico’s CARC supercomputer, with each calibration run on a CPU equipped with 2 GB of RAM.

Below we briefly summarize the results of the calibration:

1. When *γ* = 1, the down-sampled ecosystems become almost indistinguishable from the back-tracked ecosystem (see Fig. S6(a)). This is a result of the self-similarity predicted by the analytics (see Section *Analytics of the model*). This implies that there should be little difference between (*θ*^⋆^, *γ*^⋆^) and (*θ*_red_, *γ*_red_). This is verified in Fig. S6(e)–(h).
2. When *γ >* 1, the down-sampled ecosystems differ from the back-tracked ecosystems in two respects (see Fig. S6(b)): (i) the diversity of the system is underrepresented; and (ii) the abundance of the most abundant species is overestimated. For *γ >* 1, this indicates that both *θ*_red_ and *γ*_red_ are likely to be overestimated. This is verified in Fig. S6(h).
3. When *γ <* 1, the down-sampled ecosystems differ from the back-tracked ecosystems in two respects (see Fig. S6(c)): (i) the diversity is overrepresented; and (ii) the abundances of the most abundant species are underrepresented. For *γ <* 1, this indicates that both *θ*_red_ and *γ*_red_ are likely to be underestimated.
4. We find that the *γ*^⋆^ is the biggest indicator of the parameter drift in the calibrations of the down-sampled ecosystems. This is verified in Fig. S6(d) and (f), wherein the choice of *γ*^⋆^ explains most of the variance in the drift of both *θ*_red_ and *γ*_red_.
5. Drift in *θ*_red_ is found to fall into two categories: (i) When *γ* ≈ 1, *θ*_red_ is generally a slight underestimate of the true value; and (ii) When *γ* ⪆ 1, *θ*_red_ is generally an over-estimate of the true value. This is shown in Fig. S6(h).

This robustness analysis carries two important implications for the results of the model calibration conducted in the main text. The most important finding is that in the regime of linear preferential attachment, the down-sampled ecosystems are very good approximations of the back-tracked ecosystems, *implying that predictions of mechanistic model parameters should be good for sparse metagenomic data around γ* ≈ 1. This finding is convenient for the ecosystems studied herein, unlike other systems such as federal agencies and protein abundances [10]— which admit sublinear mechanisms of preferential attachment—wherein correspondence between down-sampled and back-tracked data will not hold in the general case. For systems that are not ecosystems, this implies that having access to sparse data is not enough for accurate mechanistic inference, and that procedures must be developed to deal with sparse data in those cases. A secondary point is that where super-linear preferential attachment is inferred from the metagenomic data, the underlying real ecosystem will admit super-linear preferential attachment but with a lesser strength *γ*^⋆^ *< γ*_red_.

#### Glossary of definitions and notation for robustness analysis

**Simulated ecosystem:** A simulated ecosystem of size *N* = 10^6^, with parameters *θ*^⋆^, *γ*^⋆^, and *p*_0_ = 1.

**Down-sampled ecosystem:** A limited sample of the simulated ecosystem of size *N*_red_ = 10^5^, whose species composition is sampled proportionately to the abundance of a species in the simulated ecosystem.

**Back-tracked ecosystem:** Similar to the simulated ecosystem, but where the species abundances are taken at size *N* = *N*_red_ = 10^5^. In general, this is distinct from the down-sampled ecosystem.

**Parameter drift:** Differences that arise from the true parameters describing a simulated ecosystem, and the inferred parameters from the down-sampled system.

***θ***^⋆^: The true value of *θ* for the simulated ecosystem of size *N* = 10^6^.

***γ***^⋆^: The true value of *γ* for the simulated ecosystem of size *N* = 10^6^.

***θ***_**red**_: The calibrated value of *θ* for the down-sampled ecosystem of size *N*_red_ = 10^5^.

***γ***_**red**_: The calibrated value of *γ* for the down-sampled ecosystem of size *N*_red_ = 10^5^.

## Supplementary Information

**FIG. S1:**
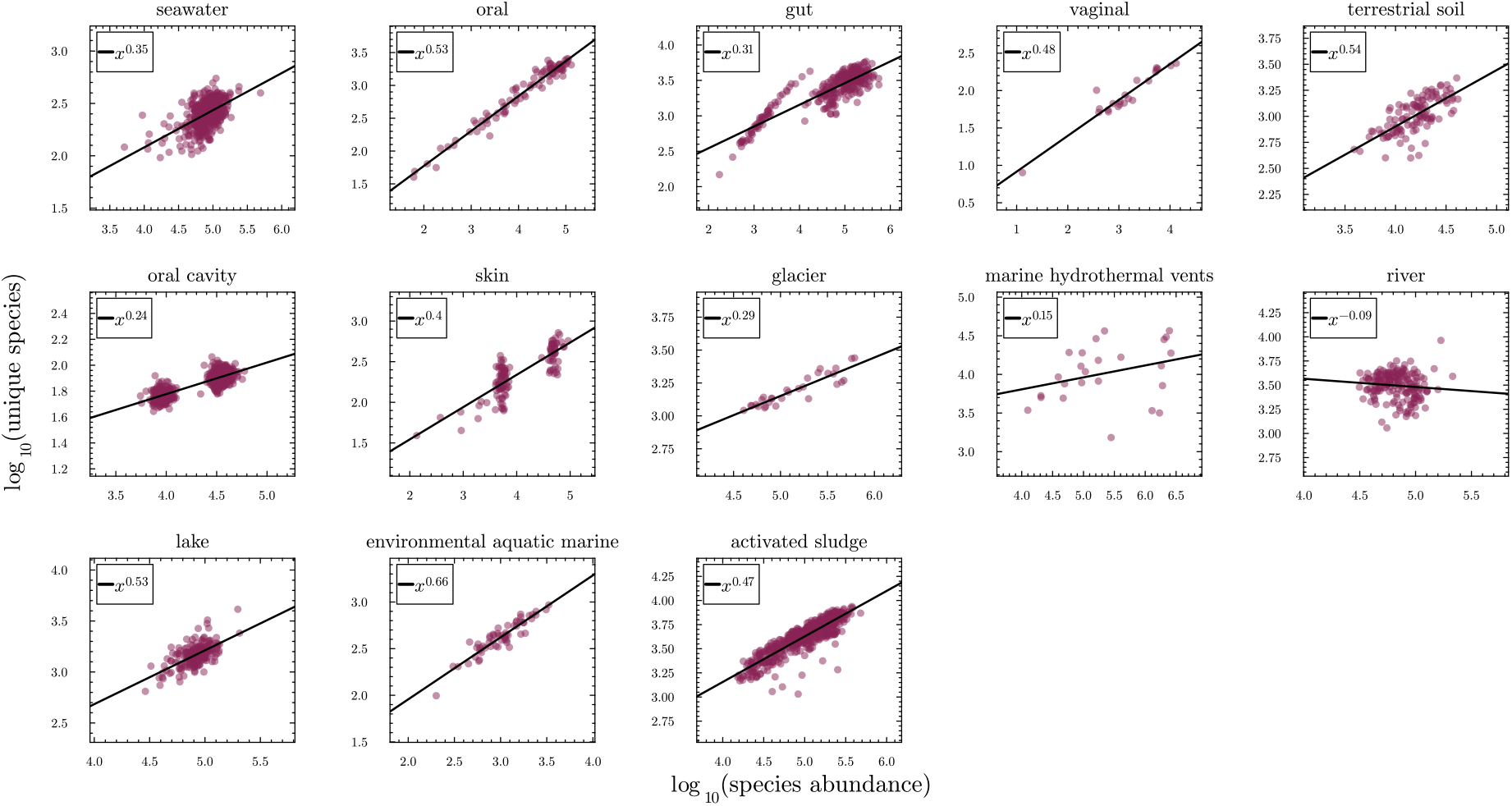
Heaps’ law across microbial ecosystems in distinct environments. Each point corresponds to a unique sample in a given environment. Fits to *D*(*N*) = *D*_0_*N* ^*β*^ were performed in Julia (for details see Table I). Exponents significantly different from zero range from 0.24 to 0.66. Two environments—*marine hydrothermal vents* and *river* —yielded Heaps’ exponents not significantly different from zero and were excluded from subsequent analyses.

**TABLE S1:**
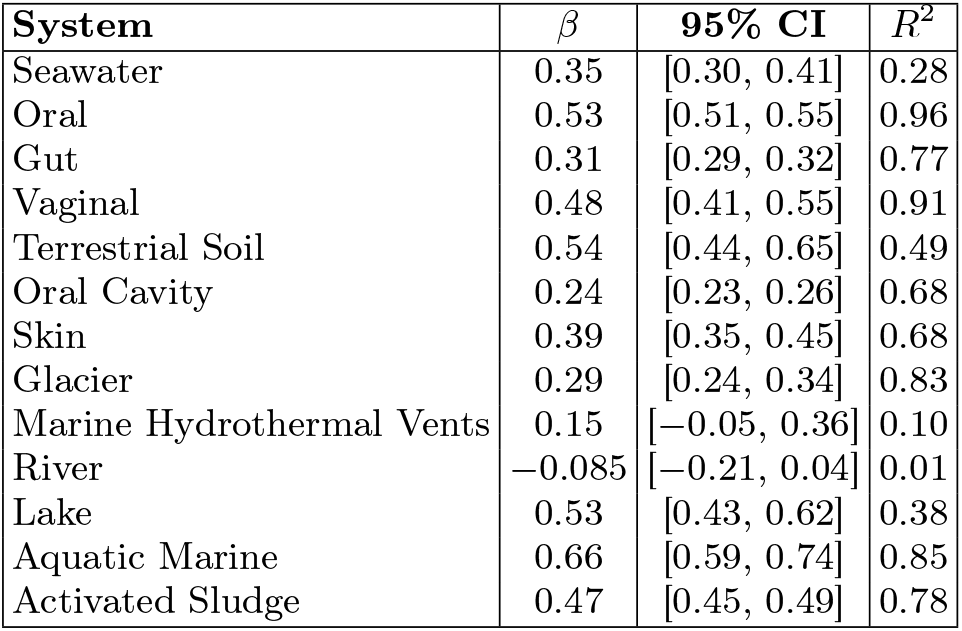
Heaps’ law fits across all environments from the dataset of ref. [1]. A linear model was fit to log *D*(*N*) = *α* + *β* log *N* using Julia’s GLM package. *R*^2^ values are evaluated in log-log space. The *marine hydrothermal vents* and *river* environments were excluded from model calibration due to non-significant Heaps’ exponents.

**FIG. S2:**
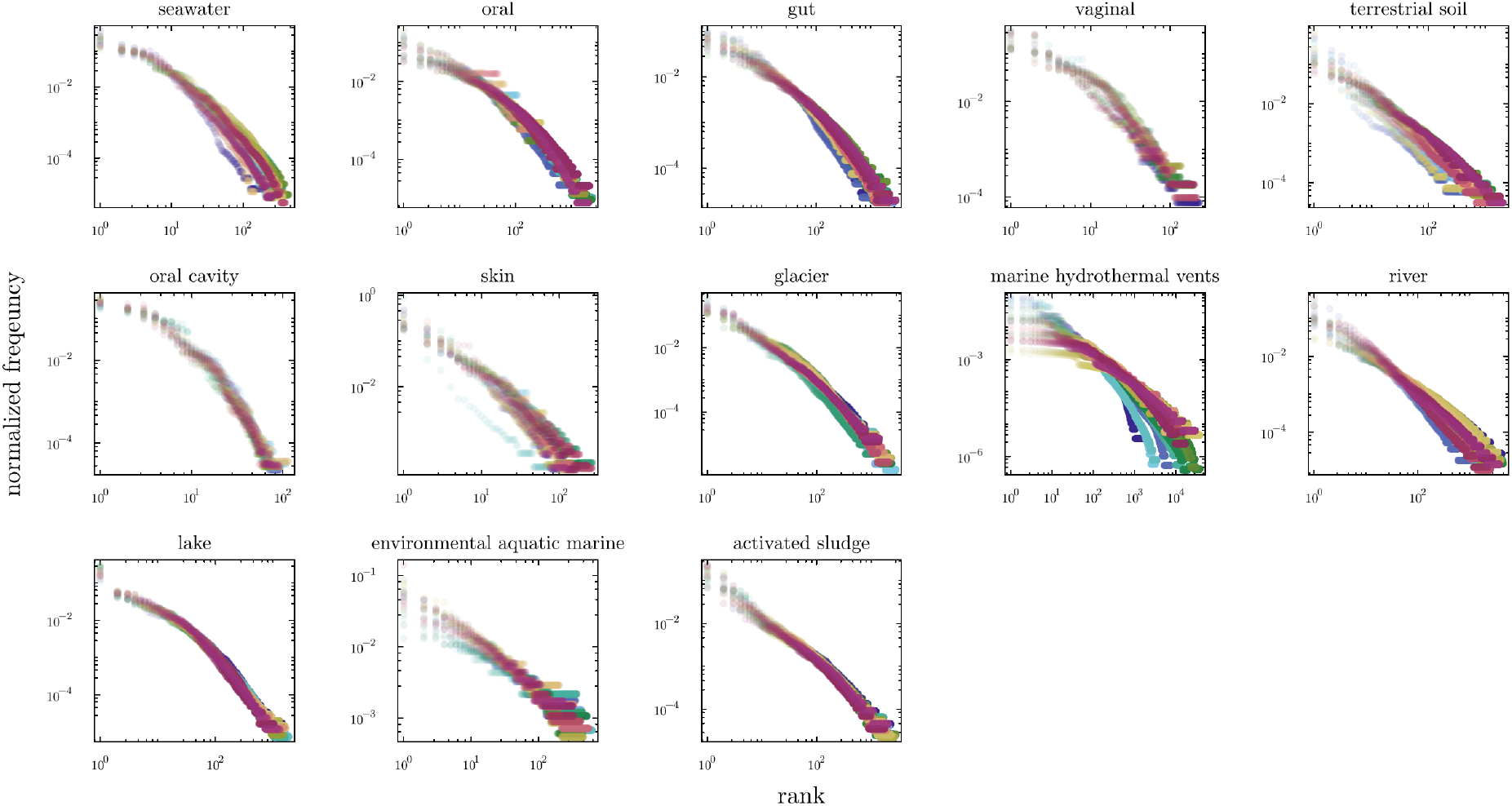
Overlapping rank-frequency distributions for 20 samples per environment. Each curve corresponds to a distinct sample, coloured by environment. Environments such as *oral, oral cavity*, and *activated sludge* display striking self-similar collapse across samples of different sizes—a direct visual signature of *γ* ≈ 1 (see *Online Methods, Analytics of the model*).

**FIG. S3:**
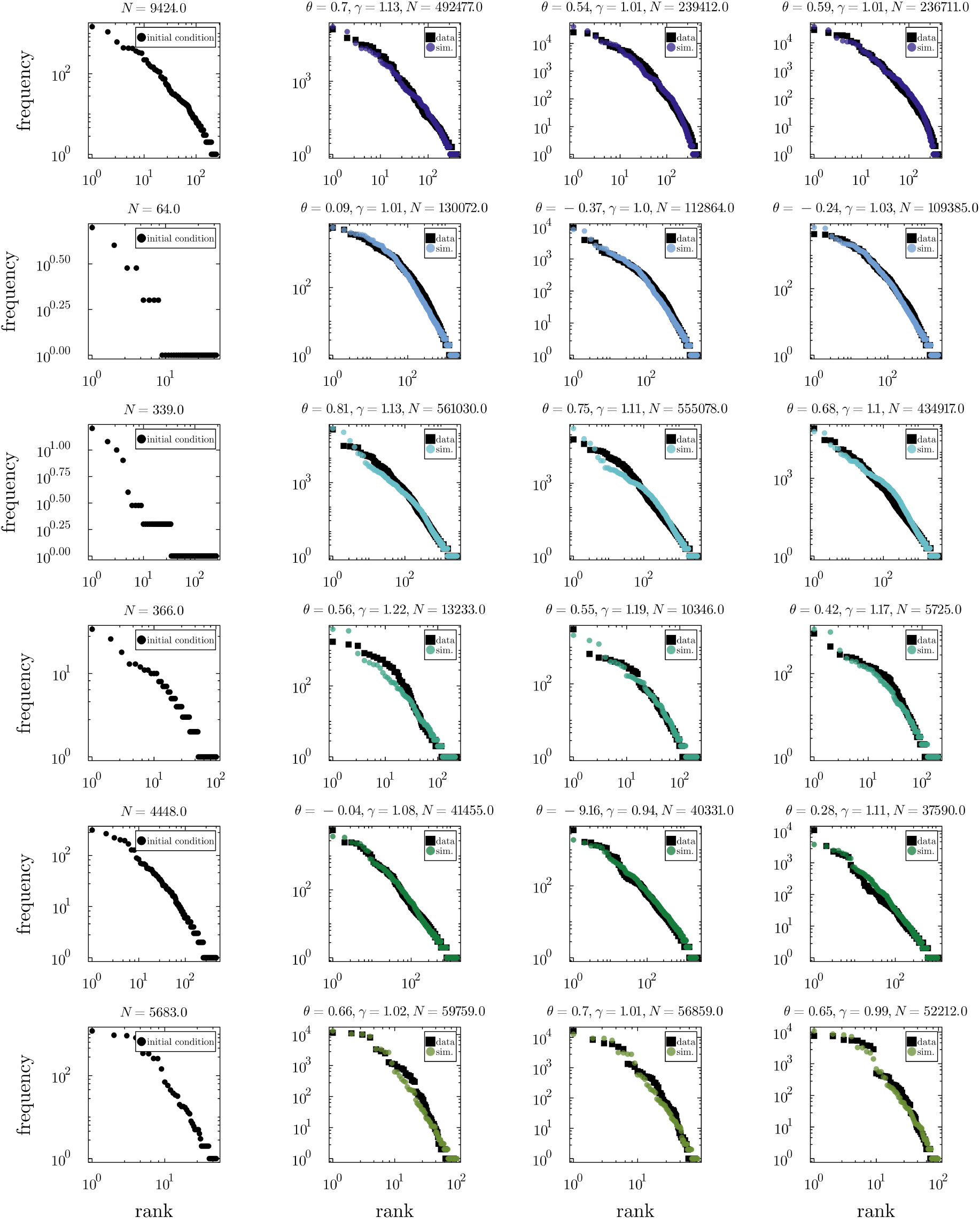
Model calibration results: seawater, oral, gut, vaginal, terrestrial soil, and oral cavity. Column 1: initial condition used for calibration (second-smallest sample in each environment). Columns 2–4: model fits to the three largest ecosystems in each environment. Plot titles report fitted parameters *θ* and *γ* and ecosystem size (*N*). Black points: empirical rank-frequency data; coloured dots: simulated data (colour scheme matches Fig. 2).

**FIG. S4:**
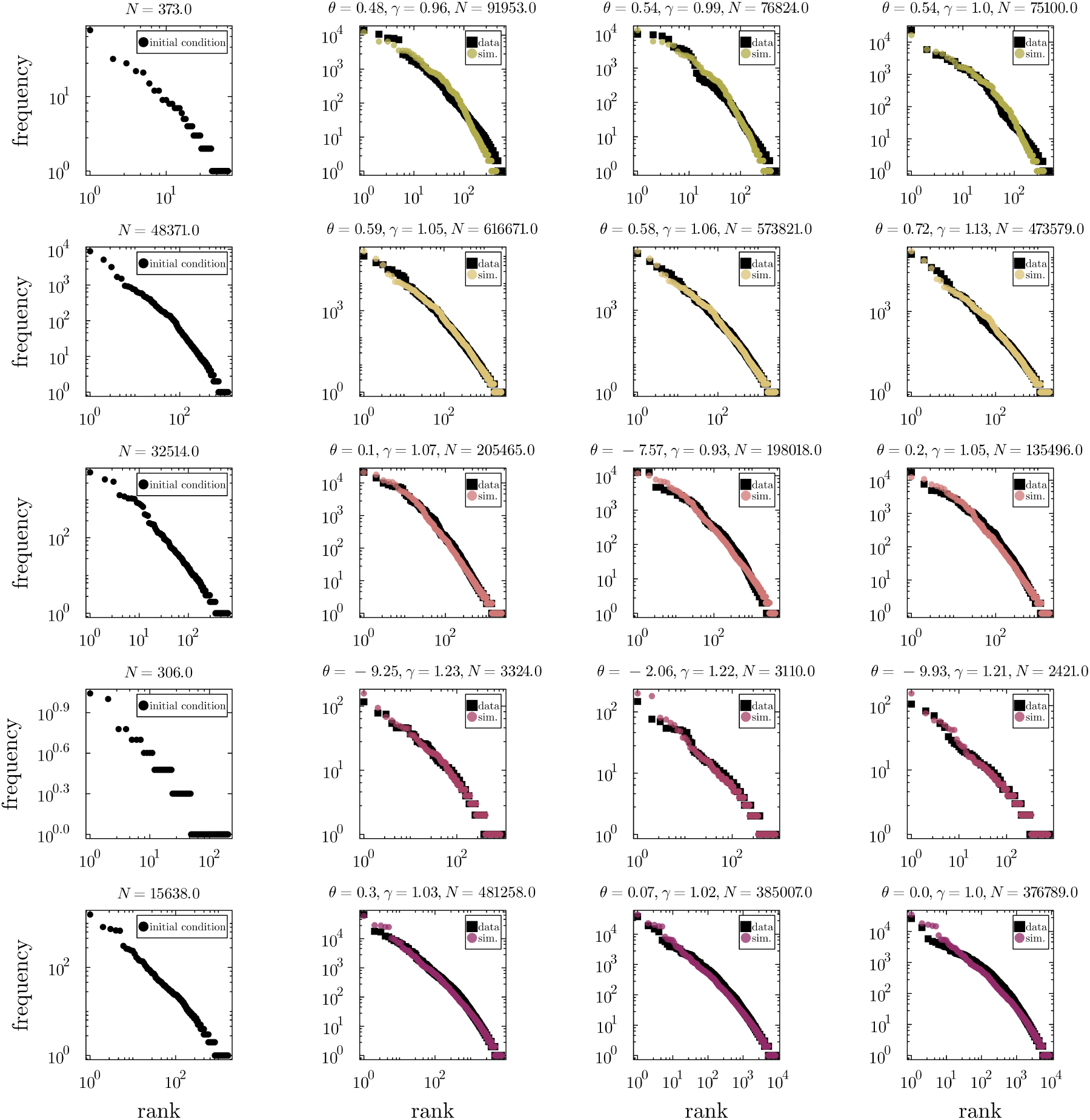
Model calibration results: skin, glacier, lake, environmental aquatic marine, and activated sludge. Layout and conventions as in Fig. S3.

**FIG. S5:**
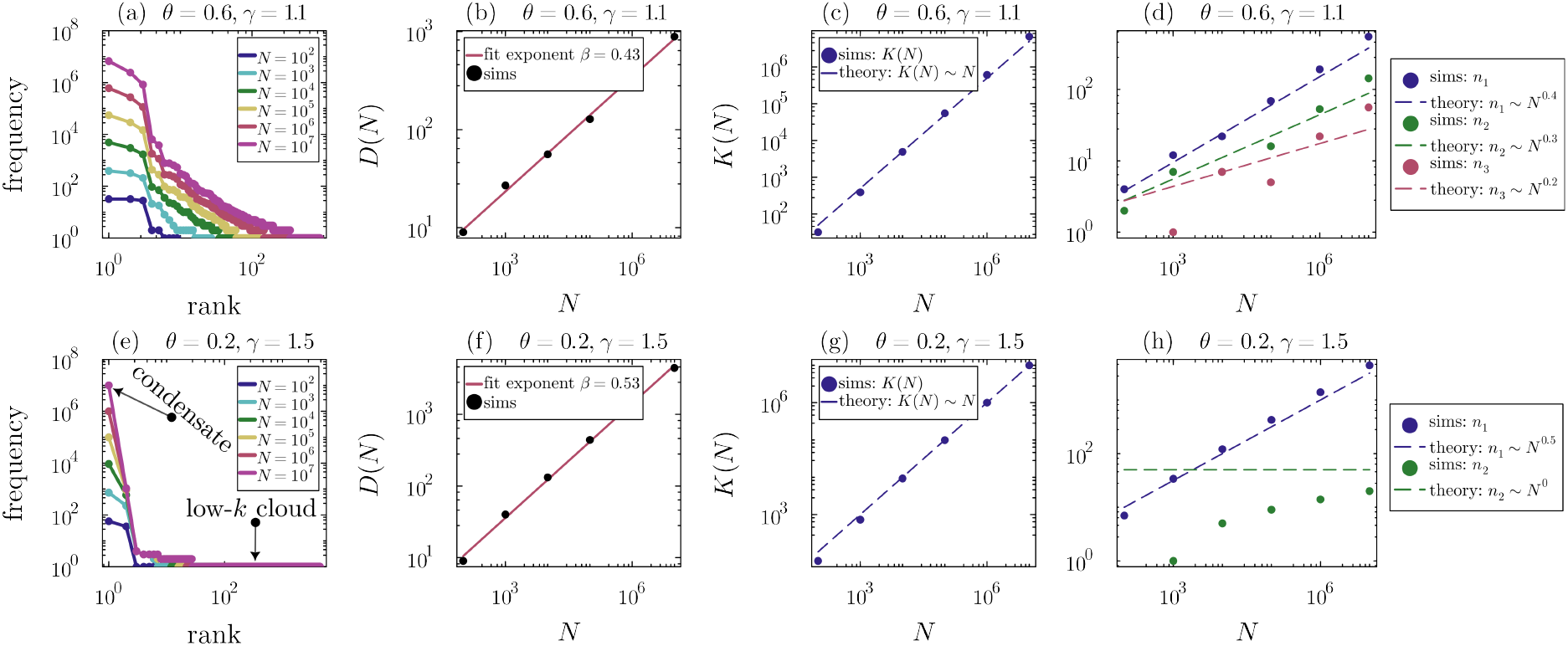
Verification of analytic results for the super-linear preferential attachment regime. (*γ >* 1**), shown for** *N* ∈ {10^2^, 10^3^, 10^4^, 10^5^, 10^6^, 10^7^}. Row 1: *θ* = 0.6, *γ* = 1.1; Row 2: *θ* = 0.2, *γ* = 1.5. Initial condition: single organism, single species. (a, e) Rank-frequency distributions showing condensate formation and a low-*k* cloud. (b, f) Diversity scaling: predicted exponents 0.4 and 0.5; observed 0.43 and 0.53. (c, g) Condensate size scales proportionally with *N* . (d, h) Scaling of *n*_*k*_ with *N* ; dashed lines show analytic predictions. Agreement is good, except for *n*_2_ in (h) where the asymptotic regime has not been reached even at *N* = 10^7^.

**FIG. S6:**
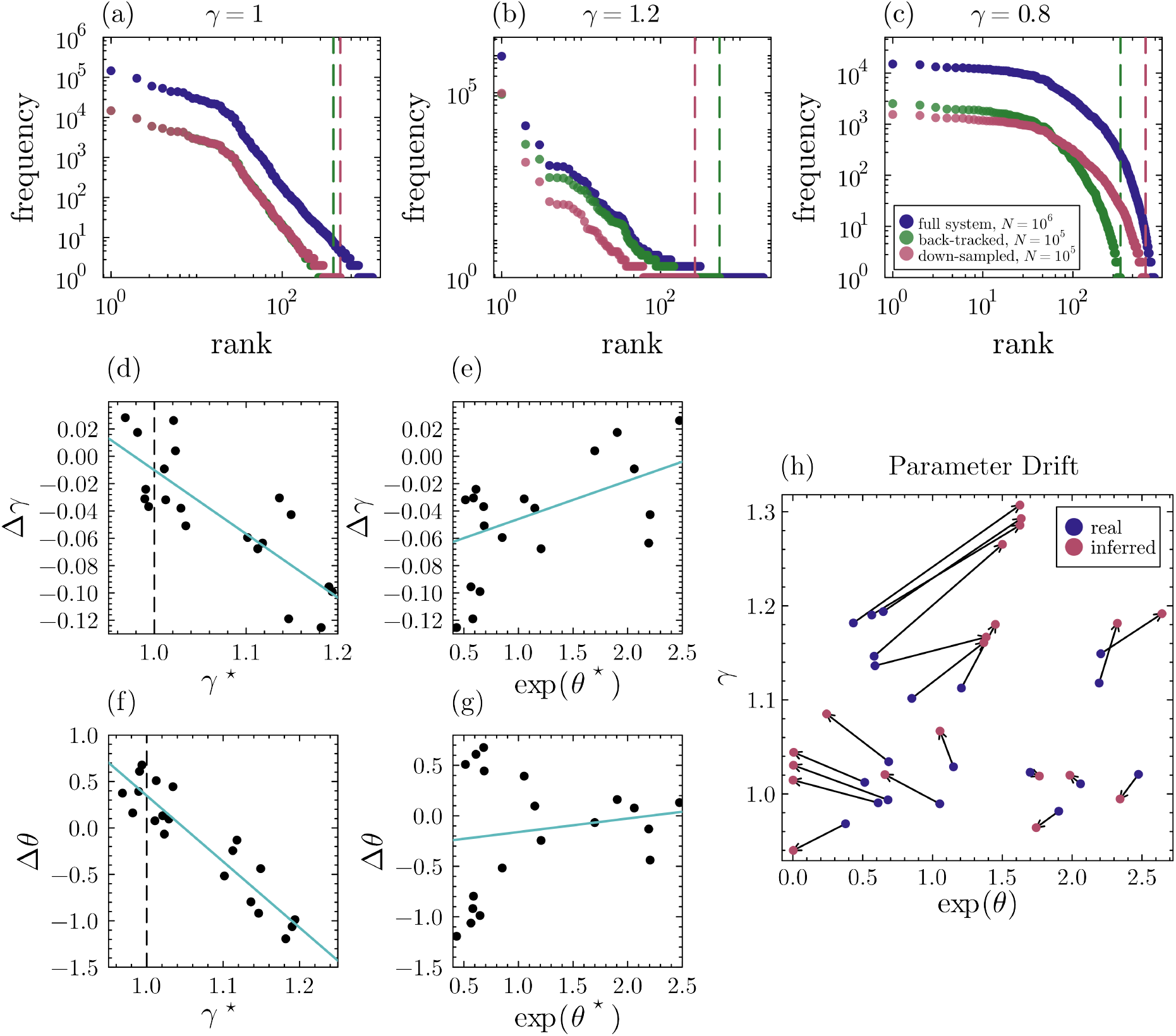
Robustness analysis of model calibration on down-sampled data. (a)–(c) For *θ* = *θ*_med_ = 0.42, ecosystems of size *N* = 10^6^ are compared after downsampling to *N*_red_ = 10^5^ (solid) vs. back-tracking to *N* = 10^5^ (dashed). (a) *γ* = 1: downsampled ≈ back-tracked. (b) *γ >* 1: diversity underestimated, dominant species overestimated. (c) *γ <* 1: opposite biases. (d)–(h) 20 ecosystems with *γ*^⋆^ ∈ [0.95, 1.2], *θ*^⋆^ ∈ [− 1, 1] downsampled and re-calibrated. (d, f) *γ*^⋆^ explains most drift: *R*^2^ = 0.67 for Δ*θ, R*^2^ = 0.86 for Δ*γ*. (e, g) *θ*^⋆^ has little predictive power. (h) True vs. inferred parameters: real and inferred pairs are closest near *γ* ≈ 1; the more super-linear *γ*^⋆^, the more super-linear the inferred *γ*_red_.

